# Human reproductive system microbiomes exhibited significantly different heterogeneity scaling with gut microbiome, but the intra-system scaling is invariant

**DOI:** 10.1101/680355

**Authors:** Sam Ma

## Abstract

Maintaining sexual reproduction in a highly competitive world is still one of the major mysteries of biology given the apparently high efficiency of asexual reproduction. Co-evolutionary theories such as the Red Queen hypothesis would suggest that the microbiomes in human reproductive systems, specifically the microbiomes contained in semen and vaginal fluids, should reach some level of homogeneity thanks to arguably the most conspicuous microbiome transmission between two sexes. The long-term sexual coevolution should favor the dynamic homogeneity or stability, which should also be beneficial for sexual reproduction such as sperm survival or fertilization on physiological/ecological time scale. We present a piece of quantitative evidence in the form of microbial community spatial heterogeneity to support the stability notion by analyzing three big datasets of the human vaginal, semen and gut microbiome. Methodologically, we applied a recent community-level extension to the classic Taylor’s power law (Taylor 1961, 1988: *Nature*), which reached the rare status of ecological law and has found applications beyond biology. The power law analysis revealed that human vaginal and semen microbiomes exhibited the same scaling parameter size in their community spatial (inter-individual) heterogeneities, while both exhibited significantly different heterogeneity scaling parameter with the human gut microbiome. Both ecological and evolutionary theories, such as hologenome/holobiont and Red Queen, even first principle, would predict that microbiome transmissions between two sexes should have homogenizing effects on the composition and stability of the microbiomes in human reproductive systems.

**Importance:** Maintaining sexual reproduction in a highly competitive world is still one of the major mysteries of biology given the apparently high efficiency of asexual reproduction. Co-evolutionary theories such as the Red-Queen hypothesis would suggest that the microbiomes in human reproductive systems, specifically the microbiomes contained in semen and vaginal fluids, should reach some level of homogeneity thanks to arguably the most conspicuous microbiome transmission between two sexes. The long-term sexual co-evolution should favor the dynamic homogeneity or stability, which should also be beneficial for sexual reproduction such as sperm survival or fertilization on physiological/ecological time scale. We present a piece of quantitative evidence in the form of microbial community spatial heterogeneity to support the stability notion by analyzing three big datasets of the human vaginal, semen and gut microbiome. Both ecological and evolutionary theories would predict that microbiome transmissions between two sexes should have homogenizing effects in human reproductive systems.

## Introduction

*Heterogeneity* is a concept studied in many fields of biology and ecology. In genetics and evolutionary biology, the importance of heterogeneity has been recognized and studied extensively since the time of Darwin (1876) in the areas of heterosis (hybrid vigor), inbreeding and genetic deterioration, based on the theory of population bottleneck that shrinking of the choice of gene variants and of potential cooperation among different gene types limits the capabilities of the restricted organism (Birchler *et al.* 2006, Wikipedia). In ecological literature, the term heterogeneity is often used informally to support the description and characterization of several similar concepts. Its interpretations are often context-dependent, and may be slightly different from its dictionary explanation—the quality or state of being diverse in character or content. Terms such as *population-, community-, ecosystem-, and landscape-heterogeneity*, are frequently used, but often not precisely defined. In the present article, we investigate the scale at the community scale. At the community level, we use heterogeneity to refer to the uneven or heterogeneous nature of species abundances among different species within a community and/or between communities, which can be quantitatively measured with an extension to the classic Taylor’s power law (Ma 2015, Li & Ma 2019). Taylor’s power law has been extensively investigated both theoretically and practically and found applications in many fields beyond its original domain of population ecology (Taylor 1961, 1984, 2007, Taylor & Taylor 1977, Taylor et al. 1983, 1988, Cohen et al. 2012, 2015; Eisler et al. 2008, Stumpf & Porter 2012; Giometto et al 2015; Ma 2015, Oh et al. 2016; Tippett & Cohen (2016); Plank & Pitchford 2017; Reuman et al. 2017; Kalinin et al. 2018).

In the present study, we comparatively investigate the heterogeneity of human microbiomes from three key habitats, *i.e*., gut, vaginal and seminal fluid. We use, to the best of our knowledge, the largest 16s-rRNA sequencing datasets in their respective sites. The objective is to determine whether there is *homogeneity* (*i.e.*, same level of heterogeneity) between the human vaginal microbiome and semen microbiome. We further compare both vaginal and semen microbiomes with a third type, arguably isolated from the both, the gut microbiome to highlight our focal objective. We were motivated to discuss possible ecological, evolutionary and reproductive implications from the comparisons.

Ecologically, the community spatial heterogeneity (CSH) is an extremely important property both theoretically and practically. For example, in the case of human microbiome, within host or intra-body microbiome heterogeneity among major microbiome habitats (including gut, skin, oral, vaginal, and lung) and inter-host (inter-subject) heterogeneity were designated as one of the primary aims of the US-NIH HMP (human microbiome project) (HMP Consortium 2012). The inter-subject heterogeneity is also a core research topic in the biogeography of human microbiome, which investigates the spatial distribution of human microbiome diversity (*e.g*., Hanson *et al.* 2012, Ma 2019). Furthermore, the heterogeneity and diversity are closely related with each other, but each with its own unique advantages in characterizing the ecological community (Ma & Ellison 2019, Ma *et al.* 2019, Li & Ma 2019).

Evolutionarily, the recently emerging *hologenome* theory of evolution recognizes that the individual animal or plant as a community or a holobiont—the host plus all of its symbiotic microbes. The theory stipulate that the variations in the hologenome—a collective genome of the holobiont can be transmitted between generations with reasonable fidelity, and are subject to evolutionarily changes caused by selection and drift (Rosenberg *et al.* 2009, Rosenberg & Zilber-Rosenberg 2018). Also according to the theory, genetic variation in the hologenome can be due to changes in the host genome as well as to changes in the microbiome, such as new acquisitions of microbes, horizontal gene transfers, and changes in microbial species abundance within hosts. Some consider the hologenome theory contains Lamarckian aspects within a Darwinian framework, accentuating both cooperation and competition within the holobiont and with other holobionts (Rosenberg *et al.* 2009, Rosenberg & Zilber-Rosenberg 2018). For example, gut microbiome was suggested to play an important role in speciation (Brucker 2013). Besides recent hologenome theory, the classic Red Queen hypotheses for explaining sexual selection and host/parasite evolutions may also be applicable to the evolution of host/microbiome co-evolution (*e.g.*, Papkou *et al.* 2018). In reproductive biology, the role of microbial symbionts in mediating reproductive isolation was extensively investigated with *Drosophila*, but without reaching a consensus (Schneider *et al.* 2019, Leftwich *et al.* 2017, Shapiro 2017). Although few similar studies have been performed in the human microbiome (Hou *et al.* 2015, Weng *et al.* 2016), the implication of microbiome in human reproductive biology cannot be excluded. Given the potentially significant ecological and evolutionary importance of the heterogeneity, Taylor’s classic power law (Taylor 1961) and its extensions (Ma 2015) offer an ideal tool to conduct our comparative analyses because its parameters (*b*) is a species or microbiome-type specific characteristic determined by the evolutionary process.

## Material and Methods

### Datasets of human gut, vaginal and semen microbiomes

We selected three large datasets of the human vaginal, gut, and semen microbiome studies with 16S-rRNA sequencing technology, as briefly introduced in Table 1. The primary considerations for selecting these three datasets include their exceptional sample size in their respective sites (vaginal, gut and semen), as well as their well-designed, high-quality sequencing operations and consequent bioinformatics analysis for generating the OTU (operational taxonomic unit) tables.

**Table 1.** Basic statistics of the three datasets: HVM (human vaginal microbiome), AGP (American gut microbiome project), and human semen microbiome

| Dataset | Number of Samples | 16S-rRNA Reads per Sample | OTU Numbers (Total) | Reference |
| --- | --- | --- | --- | --- |
| HVM | 1076 | 14,585 | 14355 | Doyle *et al.* (2018) |
| AGP | 1473 | 23,633 | 22743 | AGP:(http://americangut.org/) |
| Semen | 96 | 9,770 | 7119 | Weng *et al.* 2014 |

### Taylor’s power law and its extensions to community ecology

Taylor’s power law (Taylor 1961, 1984, 2007, Taylor & Taylor 1977, Taylor *et al.* 1983, 1988) is one of the classic mathematical models that have reached the rare status of the ecological law. It has been validated by hundreds, if not thousands of field observations in macro-ecology of plants and animals, and its theoretical implications and practical applications have extended well beyond ecology and biology, reaching fields such as epidemiology, natural catastrophe prediction, human migration, financing, and computational science (Taylor 1961, 1984, 2007, Taylor & Taylor 1977, Taylor et al. 1983, 1988, Cohen et al. 2012, 2015; Eisler *et al.* 2008, Stumpf & Porter 2012; Giometto et al 2015; Ma 2012, 2013, 2015, Oh *et al.* 2016, Tippett & Cohen (2016), Plank & Pitchford 2017, Reuman *et al.* 2017, Kalinin *et al.* 2018). In its original form, Taylor’s power law describes the relationship between population variance (*V*) and population mean (abundance) (*m*) in the following power function:

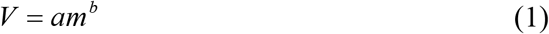

where parameter *a* is primarily influenced by the sampling scheme and environmental factors and is of limited ecological implications, and parameter *b* is of rich ecological and evolutionary implications. It is considered to be species-specific characteristic, shaped by a species’ evolutionary history and ecological interactions (Taylor 1961, 1981, 1984, 2007, Taylor & Taylor 1977, Taylor et al. 1983, 1988). Taylor’s power law was originally proposed and validated in population ecology, and parameter *b* is a measure of population *aggregation*, which characterizes the *spatial distribution* of a population in nature. When *b*>1, the population spatial distribution is *aggregated* (also termed clumped, contagious, heterogeneous); when *b*=1, the spatial distribution is *random*; when *b*<1, the distribution is uniform (also known as *regular*). Given the critical importance of population spatial distribution in population biology, Taylor’s power law, especially its aggregation parameter (*b*), is well regarded as one of the most important tools for investigating ecology and evolution of biological populations.

Taylor’s power law was extended to the community level (Ma 2015, Oh *et al.* 2016) for assessing and interpreting the community spatial heterogeneity (CSH) and/or community temporal stability (Oh et al. 2016). In the case of community spatial heterogeneity, the power law extension (PLE) has the following form:

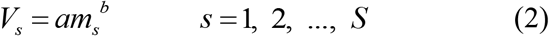

which has the same math form as the original Taylor’s power law, but with different interpretations with both the variables and parameters. In eqn. (2), *ms* is the *mean species size* (abundance) *per* species and *V*_*s*_ is corresponding variance. Parameter *a* is largely related to sampling scheme and in the case of microbiome research, it ‘absorbs’ the influence of sampling factors including the influences of sequencing platforms. This actually makes the power law advantageous because it allows for parameter *b* alone to fully capture the important ecological and evolutionary characteristic of ecological community in terms of their *spatial heterogeneity*.

The PLE parameter (*b*) offers a powerful tool to assess and interpret the community spatial heterogeneity and/or temporal stability (*e.g*., Oh *et al.* 2016). When *b*>1, community spatial heterogeneity is aggregated or asymmetrical; when *b*=1, community spatial heterogeneity is random, when *b*<1, community spatial heterogeneity is regular or uniform.

We fit the datasets of HVM (human vaginal microbiome), AGP (American gut project) and human semen microbiome (*see* Table 1) to the PLE [eqn. (2)], respectively, with an objective to compare their ecological/evolutionary characteristics, in terms of their community spatial heterogeneity as measured with PLE parameter (*b*). We utilize the randomization (permutation) test to statistically compare the PLE parameters of human gut, vaginal and semen microbiomes, as explained below.

### The randomization test procedures

To compare the power law parameters of HVM and AGP datasets, we conducted randomization (permutation) tests with the following procedures:

i. Fit the PLE to HVM and AGP datasets, respectively and obtain their respective power law parameters (as listed in Table 2); further compute the absolute differences (|*D*|) in their respective parameters.
ii. Randomly mix the samples from HVM and AGP datasets, and obtain a single pooled dataset of 2549 samples (1076HVM +1473AGP); divide the single pooled dataset into two groups, one group with 1076 samples and another group with 1473 samples, and designate them as simulated HVM and AGP dataset, respectively.
iii. Fit the PLE to each group from step *(ii)*, respectively and obtain the differences (|*D′*|) of their respective parameters.
iv. Repeat steps *(ii)* and *(iii)* for 1000 times, and obtain 1000 of *D′*-values; count the times (*n*) that satisfy |*D′*|≥|*D*| and compute a pseudo *p*-value, *i.e*., *p*=*n*/1000 for randomization test. If *p*<0.05, there is a significant difference between HVM and AGP in their respective power law parameter; otherwise, there is not.

**Table 2.** The parameters of PLE for measuring the community spatial heterogeneity of human vaginal, semen and gut microbiomes* *See Tables S1-S3 in the OSI (online supplementary information) for the PLE parameters obtained from the randomization tests for determining the differences in the heterogeneity parameters among the three microbiome types.

| Microbiome | $b$ | $\ln(a)$ | CHCD | $r$ | $p$-value | $n$ |
| --- | --- | --- | --- | --- | --- | --- |
| HMV (human vaginal microbiome) | 2.091 | 8.061 | 0.0006 | 0.902 | 0.000 | 1076 |
| AGP (American gut project) | 1.831 | 6.636 | 0.0003 | 0.797 | 0.000 | 1473 |
| Semen (human semen microbiome) | 2.327 | 5.804 | 0.0130 | 0.927 | 0.000 | 96 |

Because the sample sizes of AGP (or HVM) *vs*. semen microbiomes are rather different (1473 vs. 96), comparing the PLE parameters built with the full datasets directly could be influenced by the apparently incommensurable sample sizes. To resolve the issue, we first randomly take 96 samples from the AGP (or HVM) and those 96 samples constitute a new AGP (or HVM) group. We then use the same randomization test procedure previously designed for comparing AGP and HVM to compare the new AGP (or HVM) with semen microbiome to compare their PLE parameters.

We further repeat the above-described randomization test for 1000 times and consequently generating 1000 *p*-values. Note that, for each time of the randomization test, a new random sampling of 96 samples is performed for AGP/HVM dataset, so that the comparisons of their power law parameters with those of semen microbiome are not influenced by the sample size. Further note that, to perform each randomization test, 1000 times of re-sampling associated with standard randomization test, as introduced previously for comparing AGP vs. HVM is again taken. In other words, in each of the 1000 randomization tests, 1000 times of re-sampling for standard randomization test, was ‘embedded in’ Finally, after obtaining the 1000 *p*-values, we count the times (*N*) that satisfy *p*>0.05, i.e., no significant difference detected with randomization test, and compute a new pseudo *p*-value, *i.e*., *p=N/1000*. If *p′*<0.05, there is significant difference between HVM and semen (or between AGP and semen) in their respective power law parameters; otherwise, there is no significant difference.

## Results and Discussion

### Power law extension (PLE) models for the human vaginal, gut and semen microbiomes

The parameters of the PLE for measuring community spatial heterogeneity of human vaginal, gut and semen microbiomes were tabulated in Table 2, and the fittings were extremely significant (*p*-value<0.001). The community spatial heterogeneity parameter (*b*) for vaginal, gut and semen microbiome was 2.091, 1.831, and 2.327, respectively. Fig 1 shows the fitted power law model on log-scale in the form of linear relationship. The *b*-values, all of which are larger than *1*, indicate that the distribution of the human microbiome in all three sites (habitats) across space (individuals) are heterogeneous or asymmetrical. In terms of the property of power law, it indicates that the heterogeneity of human microbiome in a population follows a highly skewed long-tail distribution, which means that majority of individuals have relatively low variability, but small number of individuals have disproportionally large variability, in terms of their mean species abundances across different microbial species. Furthermore, there is hardly an average Joe who can represent the population he comes from, according to the so-termed “no-average” property of the power law.

**Fig 1.**
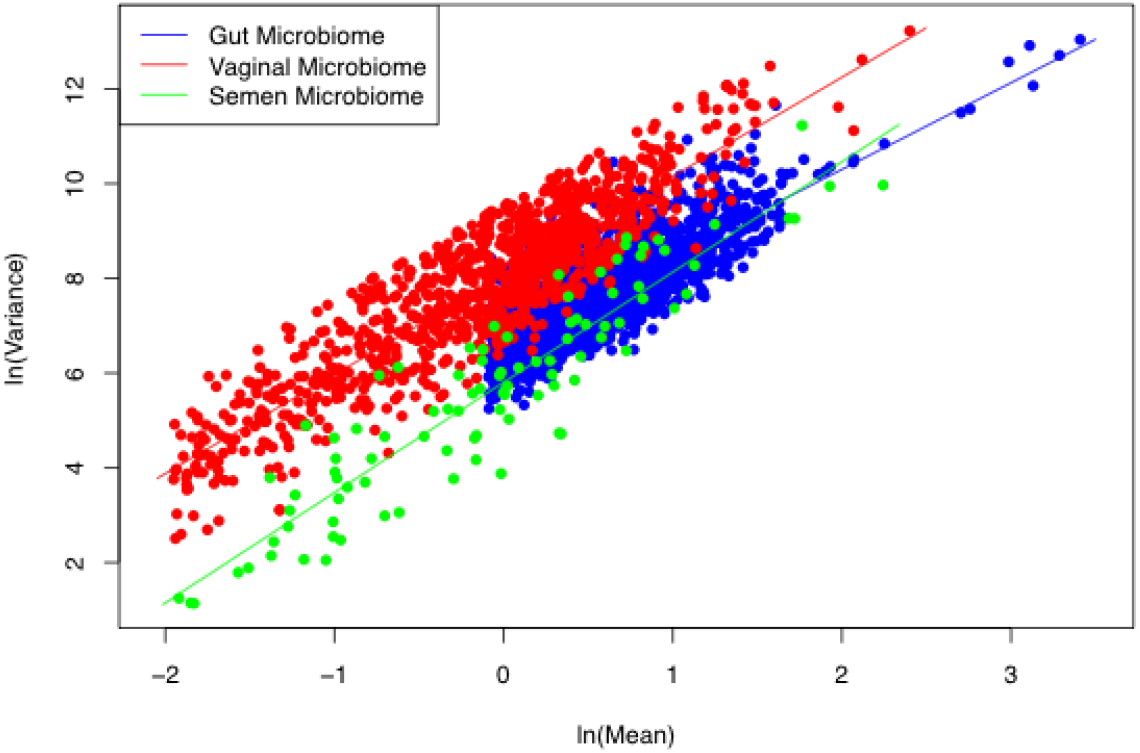
Fitting the power law extension (PLE) model for community spatial heterogeneity to HVM (human vaginal microbiome), AGP (American gut project) and semen microbiome datasets: the fitted lines for the vaginal and semen microbiomes are almost in parallel, suggesting the same slope (*b*) of both vaginal and semen microbiomes.

### Comparing the spatial heterogeneity of human vaginal, semen and gut microbiomes

The results of randomization tests (permutation tests) in Tables 3 show that the human reproductive systems (*i.e*., vaginal and semen) exhibited the same level of community spatial heterogeneity (*i.e*., *b* has no significant differences*, p>0.05*), even though we are comparing the microbiome samples from different sexes. In contrast, the human vaginal and gut microbiomes showed significant difference in the community spatial heterogeneity (*p*-value<0.001). Similarly, the human semen and gut microbiomes also showed significant difference in the community spatial heterogeneity (*p*-value<0.05). Note that our comparisons were primarily based on the power law heterogeneity parameter (*b*), but CHCD (community heterogeneity critical diversity) (Ma 2015) also followed the same trend in all three comparisons and ln(*a*) exhibited an exception in the case of semen *vs*. AGP comparison. Since parameter *a* is largely related to sampling scheme such as sequencing platforms, the exception of parameter *a* is not an issue in our analysis since we do not expect it has much ecological/evolutionary meaning. Fig 2 illustrated the triangle pattern among the human gut, vaginal and semen microbiome, in which the reproductive systems (female vaginal and male semen) exhibited the homogeneity (*i.e*., the same level of heterogeneity), but the both exhibited significantly different levels of heterogeneity with the human gut microbiome.

**Table 3.** The randomization tests for the differences in the PLE parameters among human gut, vaginal and semen microbiomes* *See Tables S1-S3 in the OSI (online supplementary information) for the detailed results of the randomization, including how the unequal sample size was dealt with.

| Microbiome | Order | Data1 | Data2 | *p*-Value |
| --- | --- | --- | --- | --- |
| HVM vs. AGP | *b* | 1.831 | 2.091 | 0.000 |
| | $\ln(a)$ | 6.636 | 8.061 | 0.000 |
|  | CHCD | 0.0003 | 0.0006 | 0.000 |
| Semen vs. HVM<br>(Sampled 1000 times) | *b* | 2.086 | 2.327 | 0.234 |
| | $\ln(a)$ | 5.904 | 5.804 | 0.786 |
|  | CHCD | 0.005 | 0.013 | 0.344 |
| Semen vs. AGP<br>(Sampled 1000 times) | *b* | 1.829 | 2.327 | 0.019 |
| | $\ln(a)$ | 6.026 | 5.804 | 0.618 |
|  | CHCD | 0.001 | 0.013 | 0.027 |

**Fig 2.**
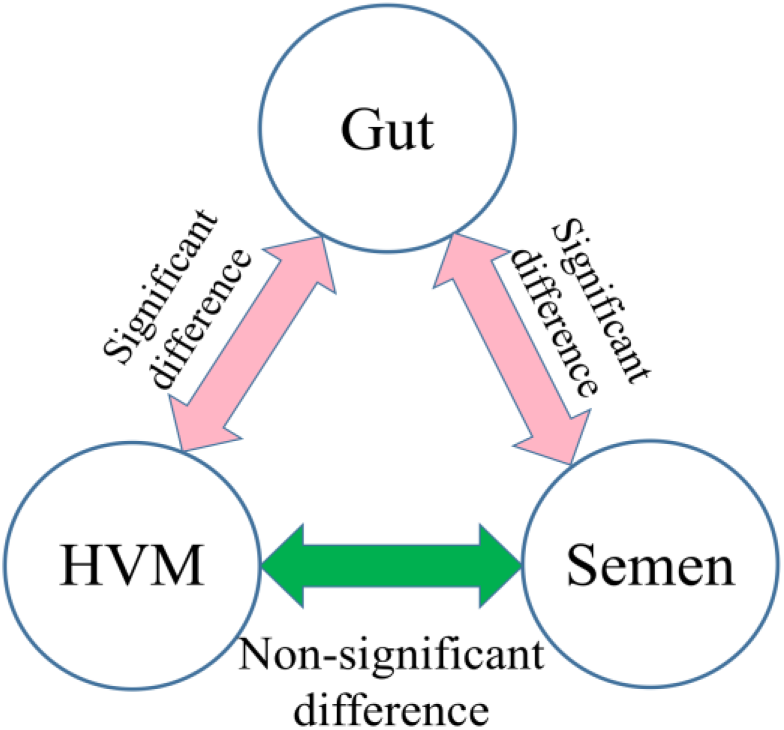
The illustration of the randomization test results presented in Table 3 (also see Tables S1-3 for the detailed results): HVM and semen microbiomes exhibited no significant difference in their community spatial (inter-subject) heterogeneity parameter, but both exhibited significant differences with the gut microbiome.

## Discussion

The results we obtained from the extended power law analysis in previous sections should come as a no surprise. First, modern human biology would expect that the human reproductive systems, including microbiomes and their hosts, should undergo co-evolutionary adaptations. The more recent *hologenome* and holobiont theory should also predict the same notions. Taylor (1961, 1984, 1986) Taylor & Taylor (1977), Taylor et al. (1983, 1988) had long been arguing that the aggregation parameter (*b*) of Taylor’s power law is a species-specific characteristic determined by a species’ evolutionary history, and recent theoretical and experimental studies have validated their early conjectures (Eisler et al. 2008; Cohen et al. 2012, 2015; Stumpf & Porter 2012; Zhang et al 2014; Giometto et al 2015; Oh et al. 2016; Tippett & Cohen (2016); Plank & Pitchford 2017; Reuman et al. 2017; Kalinin et al. 2018). The recent extensions of Taylor’s power law from population to community level should have preserved this important characteristic of parameter (*b*) (Ma 2012, 2015, Zhang et al. 2014, Oh et al. 2016, Li & Ma 2019). We argue that the difference between the concepts of *population aggregation* in original Taylor’s (1961) power law and *community heterogeneity* in the PLE (Ma 2015), to some extent, is nominal. This is because their difference is essentially single-species population *vs*. multiple-species populations. In nature, there is hardly a single species population that exists in isolation from other species, and human microbiome is no exception. What differ are the different levels or even kinds of interactions between co-specific and inter-specific individuals. But when captured by heterogeneity, both kinds (levels) of interactions are on the same metric dimension. Therefore, the nominal or apparent difference between the original power law (Taylor 1961) and the power law extension (Ma 2015) is simply a change of counting system of organisms, not unlike the relationship between binary and decimal systems in computer science. In other words, population aggregation (measured by original Taylor’s power law) and community heterogeneity (measured by the power law extension) are both evolutionary characteristics, exhibited at different scales (population *vs*. community or species *vs*. microbiome).

In the case of the human microbiome, the relationship between the microbes and their host (or our body) are so tightly connected that there have been suggestions to treat gut microbiome as a human organ. The situations between human vaginal and vaginal microbiome or between semen microbiome and seminal fluid should be similarly close. Therefore, the relationships we analyzed in this article should be the product of evolution. The dispersal (migration or transmission) occurred between microbes in human vaginal and seminal fluid is arguably the most important inter-human transmission, besides mother-baby microbiome transfer during the birth and breastfeeding. This dispersal is obviously of significant ecological and evolutionary implications. Ecologically, it is critical for shaping the structure and dynamics of the metacommunities of human microbiomes hosted by human populations, in particular, the microbes hosted by human reproductive systems. Evolutionarily, the dispersal of microbes between both sexes should have played a significant role in driving the microbiome evolution as well as their relationships with the human reproductive systems. Both ecological and evolutionary theories would predict that dispersal between two sexes should have homogenizing effects on the composition and stability of the microbiomes in human reproductive systems. Our power law analysis provides the very first piece of quantitative evidence to measure the homogenization of microbiomes within the human reproductive systems.

While dispersal should promote homogenization ecologically and evolutionarily can be justified by *first principle of dispersal physics*, what are the benefits of such homogenization in terms of reproductive fitness? We postulate that it is the *stability* of microbiome that matters for the reproductive success. Homogenization between human semen and vaginal microbiomes should be beneficial for stabilizing the microbial environments of reproductive systems. In other words, difference in heterogeneity levels between semen and vaginal would means the high *potential* for changes or instability, which may not be a benign environment for the life of sperm or for fertilization to occur. This hypothesis is certainly subject to future studies to confirm or reject. As a side note, the type-III power law extension for measuring community temporal stability (Ma 2015) has been successfully applied for skin microbiome stability (Oh *et al.* 2016). However, currently, there is not long enough time-series data of the human gut or semen microbiome available in existing literature, to conduct similar comparative analysis for the temporal version of the power law extension. We hope that future studies will fill this gap. It should certainly be interesting to compare the temporal stabilities of human gut, vaginal and semen microbiomes.

In a recent comparative study of the extended power law parameters between the hot spring microbiome and human gut microbiome, Li & Ma (2019) found that the heterogeneity scaling parameter (*b*) of hot spring microbiome is invariant with hot spring environments such as temperature and acidity (pH). However, the heterogeneity scaling parameter (*b*) of the hot spring microbiome was significantly different from that of the human gut microbiome. They used an analogy with the gravitational acceleration rates of earth and moon, which are different on earth and moon. Analogically, the human and hot spring can possess different heterogeneity scaling parameters. Of course, the gravitational acceleration on the earth or moon should be invariant or constant (despite slight differences exists on different latitudes and longitudes), just like the scaling of hot spring microbiome is invariant with temperatures or pH. Li & Ma (2019) finding echoed our previous finding in this study—the invariance of the inter-individual heterogeneity scaling of the human reproductive system microbiomes. Similar to the difference between the earth and moon in their gravitational acceleration rates, the difference in the heterogeneity scaling between the reproductive system and digestive system should come as a no surprise due to their functional differentiations.

## Author Contributions

ZS Ma designed and conducted the study, and wrote the paper.

## Data accessibility

All the datasets used in this study are available in public domain as listed in Table 1.

## Compliance with ethical standards

N/A

